# Terraces in Species Tree Inference from Gene Trees

**DOI:** 10.1101/2022.11.21.517454

**Authors:** Mursalin Habib, Kowshic Roy, Saem Hasan, Atif Hasan Rahman, Md. Shamsuzzoha Bayzid

## Abstract

A terrace in a phylogenetic tree space is a region where all trees contain the same set of subtrees, due to certain patterns of missing data among the taxa sampled, resulting in an identical optimality score for a given data set. This was first investigated in the context of phylogenetic tree estimation from sequence alignments using maximum likelihood (ML) and maximum parsimony (MP). The concept of terraces was later extended to the species tree inference problem from a collection of gene trees, where a set of equally optimal species trees was referred to as a “pseudo” species tree terrace. Pseudo terraces do not consider the topological proximity of the trees in terms of the induced subtrees resulting from certain patterns of missing data. In this study, we mathematically characterize species tree terraces and investigate the mathematical properties and conditions that lead multiple species trees to induce/display an identical set of locus-specific subtrees owing to missing data. We report that species tree terraces are agnostic to gene tree topologies and the discordance therein. Therefore, we introduce and characterize a special type of gene tree topology-aware terrace which we call “peak terrace”, and investigate conditions on the patterns of missing data that give rise to peak terraces. In addition to the theoretical and analytical results, we empirically investigated different challenges as well as various opportunities pertaining to the multiplicity of equally good species trees in terraced landscapes. Based on an extensive experimental study involving both simulated and real biological datasets, we present the prevalence of species tree terraces and the resulting ambiguity created for tree search algorithms. Remarkably, our findings indicate that the identification of terraces and the trees within them can substantially enhance the accuracy of summary methods. Furthermore, we demonstrate that reasonably accurate branch support can be computed by leveraging trees sourced from these terraces.

## 1 Introduction

Species tree estimation is frequently based on phylogenomic approaches that use multiple genes from throughout the genome. The estimation of species trees from multiple genes is necessary since true gene trees can differ from each other and from the true species tree due to various processes, including gene duplication and loss, horizontal gene transfer, and incomplete lineage sorting (ILS) [1]. A traditional approach to species tree estimation from multi-locus data is called concatenation (also known as combined analysis), where alignments are estimated for each gene and concatenated into a supermatrix, which is then used to estimate the species tree using a sequence based tree estimation technique (e.g., maximum parsimony, maximum likelihood etc.). The concatenation approach, which is agnostic to the topological differences among the gene trees, can be statistically inconsistent [2] and can return incorrect trees with high confidence [3–6].

As a result, “summary methods”, that operate by summarizing estimated gene trees and can explicitly take gene tree discordance/heterogeneity into account are becoming increasingly popular [7–18]. Fundamental to most of these summary methods is the ability to search the tree space under certain optimization criteria (e.g., maximizing pseudo likelihood score [12], maximizing quartet score [11, 18, 19], maximizing triplet score [17], minimizing deep coalescence [20]). As the size of the tree space grows exponentially with the number of taxa, finding the optimal species tree with respect to a particular optimization criterion is challenging. Moreover, the presence of local optima and multiple optimal solutions make the tree search even more complicated.

A concept related to the presence of multiple optimal solutions is called “phylogenetic terraces” – regions of the tree space where all trees have the same score due to certain patterns of missing data. Sanderson *et al*. [21] first formally investigated this and showed that when phylogenetic trees are estimated from sequence alignments using maximum likelihood (ML), multiple distinct trees can have exactly the same likelihood score due to missing data (i.e., missing genes) – a phenomenon which was referred to as terraces and was further investigated in subsequent studies [22–25]. Farah *et al*. [26] showed that similar phenomenon can arise when species trees are estimated by summarizing a collection of gene trees. They introduced the concept of “pseudo species tree terrace”, where potentially large numbers of distinct species trees may have exactly the same optimality score with respect to a set of input gene trees. There could be many reasons for multiple trees to have an identical score, but the trees in a terrace are indistinguishable in an important way: they “display” the same set of subtrees which subsequently results in identical optimality scores. For a species tree *T* and and a locus/gene tree *gt*, the locus-specific induced subtree of *T* is obtained by pruning the taxa in *T* that are missing in *gt*. Two topologically different trees *T*_1_ and *T*_2_ can induce the same locus-specific subtree due to certain patterns of missing data. This type of topological closeness in terms of identical sets of induced subtrees was not considered in pseudo species tree terraces (this is why it was called a “pseudo” terrace). Therefore, pseudo species tree terraces can arise even without the presence of missing data.

The discovery of species tree terraces has implications for summary methods that navigate through and score the trees within the tree space. Because all of the trees within a species tree terrace have the same optimization score, recognizing a terrace may help reduce computing efforts by avoiding computation time that would otherwise be spent evaluating many trees with identical scores. However, it is possible that some trees in a terrace are topologically more correct than the other ones, which was systematically analyzed and empirically demonstrated in [26]. As a result, in the presence of potentially large sets of equally optimal trees, detecting terraces and identifying relatively more reliable trees within the terraces and their neighborhoods may improve the performance of tree search algorithms. Indeed, *terrace-aware* data structures led to substantial speedup of RAxML [27, 28] and IQ-tree [29] for estimating ML trees from alignments [23].

The conditions for datasets to have phylogenetic terraces, described in [21], are general and extensible to the gene tree-species context [26, 30]. However, the combinatorial properties and mathematical characterizations of species tree terraces and the characteristics of the input gene trees and missing data patterns that lead to the presence of species tree terraces have not been elucidated in the gene tree-species tree context. In this paper, we mathematically characterize the species tree terrace and investigate various combinatorial properties of terraces. Moreover, we show that species tree terraces are not sensitive to gene tree topologies and their discordance and as a result, one set of species tree acts as a species tree terrace for an extremely large number of different sets of input gene trees. Therefore, we introduce and formalize a special type of gene tree topology-aware species tree terrace “peak terrace”, describe its importance, argue why it suffices to only look at them to understand the properties of species tree terraces in general and investigate conditions on the patterns of missing data and taxon coverage that give rise to peak terraces. In our study, we further explored, using a collection of simulated and real datasets, the presence and impact of species tree terraces and peak terraces. We showed that summary methods (e.g., ASTRAL, wQFM, etc.) may frequently estimate trees that fall within large species tree terraces with associated challenges in distinguishing trees in terms of their accuracy. In this connection, we investigated various ways to address these challenges associated with species tree terraces. We show that substantial improvements in species tree accuracy could be achieved if we can effectively leverage the trees inside a terrace. Moreover, we investigated the potential for estimating branch supports of a species tree using trees within a terraced landscape.

## 2 Preliminaries

We now define some general terminology we will use throughout this paper; other terminology will be introduced as needed. All trees *T* that follow are full binary trees with node set *V* (*T*), edge set *E*(*T*), and leaf set *L*(*T*). Let *T* be a full binary tree and let *X* ⊆ *L*(*T*). The homeomorphic subtree of *T* induced by *X*, denoted by *T* |*X*, is the unique tree obtained by restricting *T* to the leaf set *X* and then suppressing all the nodes of degree two in the resulting tree (see Figure 1). If *T* ^*′*^ is a homeomorphic subtree of *T*, then we say that *T displays T* ^*′*^. We consider the *restriction-based* approach [31–33] where an incomplete gene tree (i.e., a tree that can miss some taxa) *gt* is reconciled with a species tree *T* by taking the homeomorphic subtree *T* ^*′*^ = *T* |*L*(*gt*).

**Figure 1:**
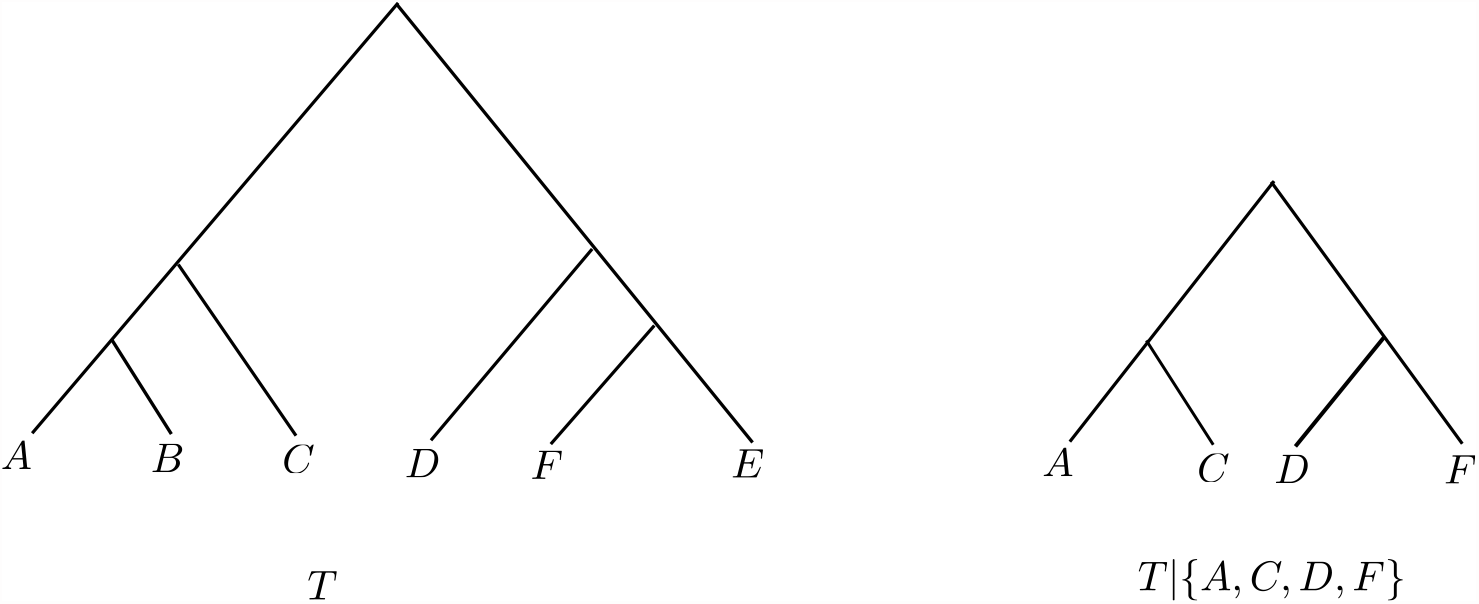
A full binary tree *T* and its homeomorphic subtree *T* |*X* where *X* = {*A, C, D, F*}

The problem of inferring species trees from multi-locus data using summary methods generally involves the following setup: we have as input a sequence *𝒢* = (*g*_1_, *g*_2_, …, *g*_*k*_) of *k* gene trees such that *L*(*g*_*i*_) ⊆ *𝒳* for each *i ∈* {1, 2, …, *k*} and we wish to find a species tree *T* that is optimal with respect to *𝒢* according to a predefined scoring function *s*_*G*_. There are many scoring functions of interest such as the extra lineage (due to deep coalescence) score [20], the pseudo-likelihood score [12], triplet score [17], and the quartet score [11]. In this article, we will focus on the quartet score, but our results are general and extensible to other optimization criteria as well. A *quartet* is a binary tree on four leaves. We denote by *ab*|*cd* the unrooted quartet tree in which the pair *a, b* is separated from the pair *c, d* by an edge. Given a binary tree *T* with at least four leaves, we denote by *Q*(*T*) the set of all quartets displayed by *T*. The *quartet score* of a species tree *T* with respect to an input sequence 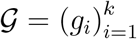 of gene trees is given by 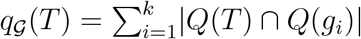.

## 3 Species tree terraces

The concept of phylogenetic terraces [21], originally described for tree estimation from sequence data using maximum likelihood, was later extended to species tree estimation from gene trees using summary approaches in [26], which showed that for a fixed sequence *𝒢* of gene trees, there can be potentially large sets of species trees with identical optimality scores. These sets of equally good species trees can arise regardless of the presence of missing data and were referred to as pseudo species terraces (see Definition 3.1).

### Definition 3.1 (Pseudo Species Tree Terrace)

*Let* 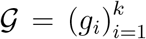 *be a sequence of gene trees and let s*_*𝒢*_(*·*) *be a scoring function. A pseudo species tree terrace is a set 𝒮 of the following form*.

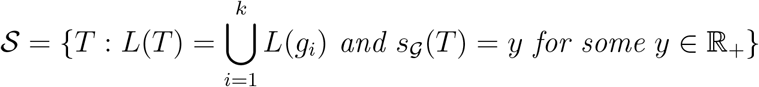

See Figure 2 for an example of a pseudo quartet terrace. Note that every tree in this pseudo terrace has the same quartet score with respect to the input sequence of gene trees. However, some of these trees have the same score due to a very specific reason – they display the same set of subtrees. In order to highlight this point, we define *species tree terraces* which are pseudo species tree terraces with an additional condition.

**Figure 2:**
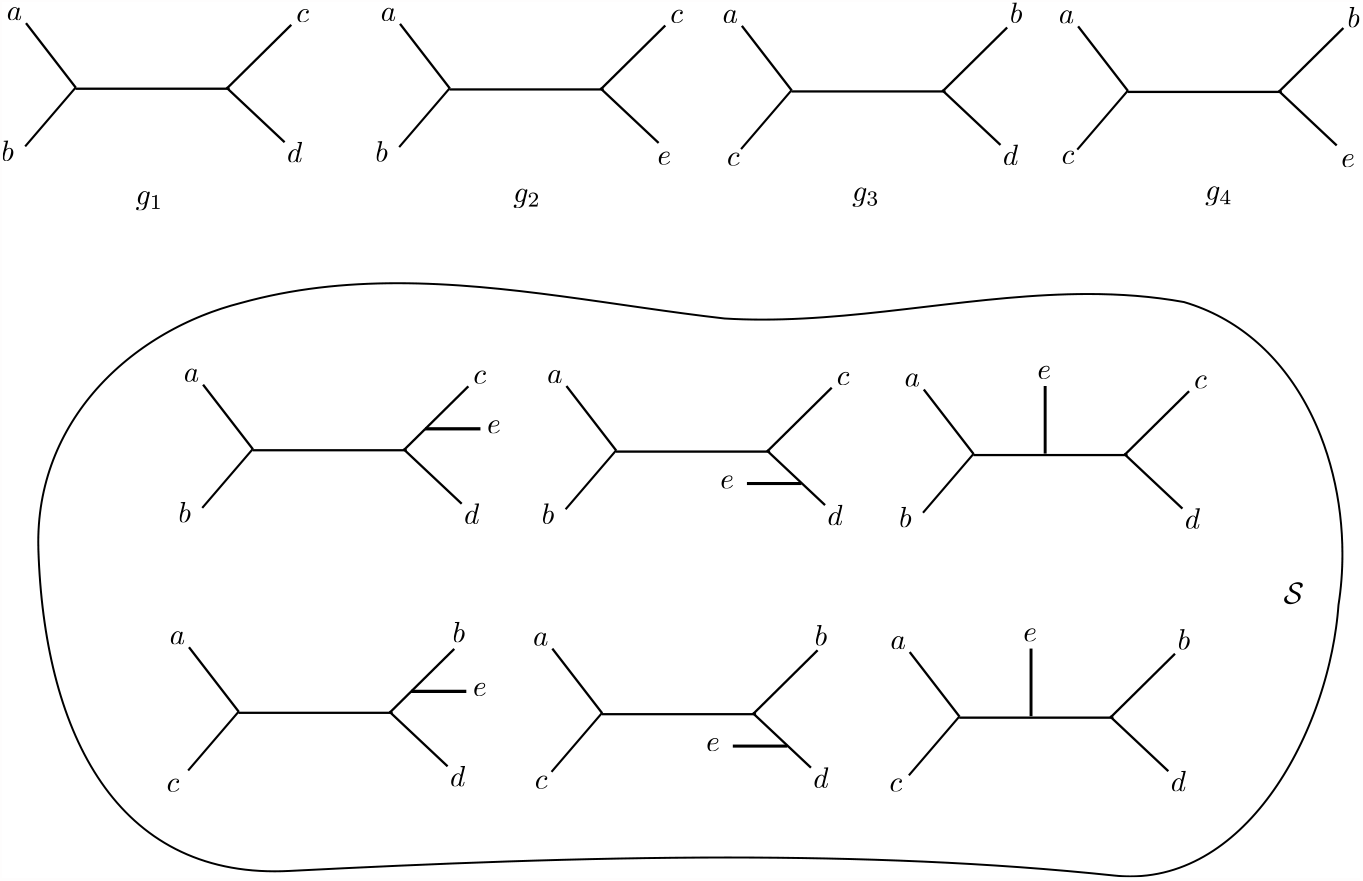
A pseudo quartet terrace *𝒮* for the sequence *𝒢* = (*g*_1_, *g*_2_, *g*_3_, *g*_4_) of gene trees. Any tree in *𝒮* has a quartet score of 2 with respect to *𝒢*.

### Definition 3.2 (Species Tree Terrace)

*Given an input sequence* 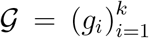 *of gene trees, two species trees T and T* ^*′*^ *are said to be in a species trees terrace if both of the following conditions hold*.

1. *T and T* ^*′*^ *reside in the same pseudo species trees terrace with respect to 𝒢, and*
2. *the sequences* 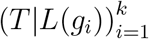 *and* 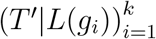 *are equal*.

In other words, trees in a species tree terrace not only have the same score with respect to *𝒢* but also display the same homeomorphic subtrees when restricted to the leaves of the input gene trees. A few comments about Definition 3.2 are in order. Firstly, most scoring functions we care about turn out to have certain common properties. Given a sequence 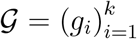 of gene trees, we call a scoring function *s*_*𝒢*_ (*·*) *additive* if the following holds for every species tree *T*.

1. There exists a function *c* such that *c*(*T, g*_*i*_) = *c* (*T* |*L*(*g*_*i*_), *g*_*i*_) for all 1 *≤ i ≤ k*, and
2. 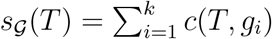.

Clearly, the quartet score *q*_*𝒢*_ (*·*) is additive with *c*(*T, g*_*i*_) being the number of quartets common to both *T* and *g*_*i*_ i.e., |*Q*(*T*) *∩ Q*(*g*_*i*_)|. The reader is encouraged to verify that other scores such as the extra-lineage score, the triplet score, the gene duplication and loss score [32, 33], and the pseudo likelihood score [12] are also additive.

It turns out that if we are dealing with only additive scoring functions, condition 1 in Definition 3.2 is redundant. In other words, we have the following.

### Proposition 3.1 (From [30])

*Let* 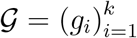 *be a sequence of gene trees and let s*_*𝒢*_ (*·*) *be an additive scoring function. If T and T* ^*′*^ *are species trees with* 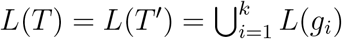 *that satisfy* 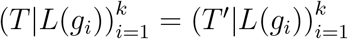, *then s*_*𝒢*_ (*T*) = *s*_*𝒢*_ (*T* ^*′*^).

One consequence of Proposition 3.1 is that if two trees are in a species tree terrace for a certain additive scoring function (say, the quartet score), then they are in a species tree terrace for *every* additive scoring function. In other words, the choice of the scoring function is irrelevant as long as it is additive. This motivates us to define an “additive version” of a species tree terrace that is independent of the choice of the scoring function.

### Definition 3.3 (Additive Species Tree Terrace)

*Given an input sequence* 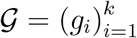 *of gene trees, two species trees T and T* ^*′*^ *with* 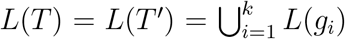 *are said to be in an additive species trees terrace if* 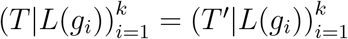.

This immediately lets us conclude the following corollary.

### Corollary 3.1

*Let T and T* ^*′*^ *be species trees residing in an additive species tree terrace with respect to a gene tree sequence* 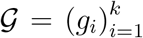. *Then T and T* ^*′*^ *have equal score with respect to any additive optimization criteria (e*.*g*., *quartet score, triplet score, extra-lineage score, and gene duplication and loss score)*.

Since our focus is the quartet score which is an additive scoring function, for the remainder of the article we use “species tree terrace” and “additive species tree terrace” interchangeably.

Another point worth noting is that Definition 3.2 does not disallow terraces of cardinality one. In fact, given 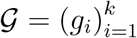, every species tree *T* with 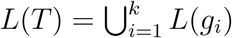 is part of some terrace (containing possibly only *T* itself). Indeed, let *T* be an arbitrary species tree with 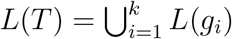 and consider the set *𝒮* = {*T* ^*′*^ : *L*(*T* ^*′*^) = *L*(*T*) and 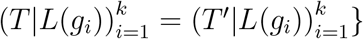. Clearly, *T* ∈ *𝒮* and therefore, |*𝒮*| *≥* 1. If |*𝒮*| *>* 1, then *𝒮* is clearly a species tree terrace. However, note that even if |*𝒮*| = 1, the conditions in Definition 3.2 are vacuously true and *𝒮* is technically a terrace. Such terraces of cardinality one are not very interesting in the context of phylogenomic analyses. Rather a much more interesting task is to investigate the conditions for which there exist more than one tree in a particular terrace.

From Definitions 3.1 and 3.2, it is clear that every pseudo species tree terrace can be partitioned into a set of species trees terraces. In other words, there may be multiple terraces imbedded in a larger pseudo terrace at the same “elevation” in the landscape of tree space. Figure 3 shows how the pseudo quartet terrace *𝒮* in Figure 2 is composed of two different quartet terraces *𝒮*_1_ and *𝒮*_2_. All of the trees in *𝒮*_1_ (and likewise in *𝒮*_2_) display identical trees when restricted to the leaf set of a particular gene tree. For example, when restricted to *L*(*g*_1_), all the trees in *𝒮*_1_ and *𝒮*_2_ display ((*a, b*), (*c, d*)) and ((*a, c*), (*b, d*)), respectively.

**Figure 3:**
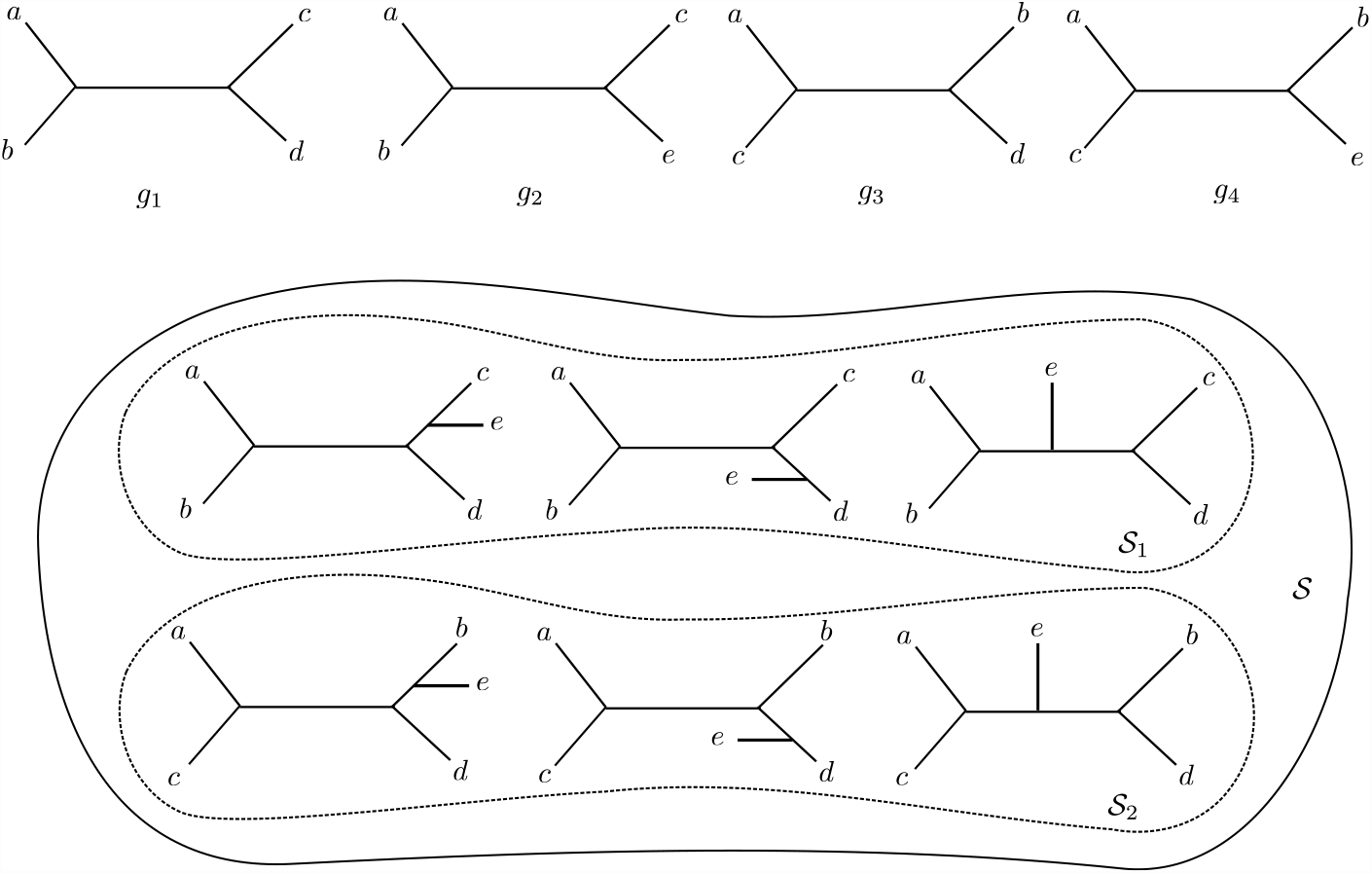
The pseudo quartet terrace of Figure 2 partitioned into two species tree quartet terraces *𝒮*_1_ and *𝒮*_2_.

Note that due to the extra condition that trees in a species tree terrace must satisfy, terraces of size greater than one can only exist if there is missing data in the input gene trees i.e. for every 1 ≤ *i* ≤ *k*, 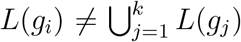. This is because if there exists *i* such that *g*_*i*_ contains data from all the taxa, then for any species tree *T, T* |*L*(*g*_*i*_) = *T*, and the only tree with leaf set 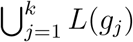 that displays *T* is *T* itself. Pseudo species tree terraces have no such restrictions: large pseudo-terraces exist even without the presence of any missing data [26]. Some lower bounds on the sizes of such large pseudo terraces can also be derived as shown below.

### Theorem 3.1

*Let* 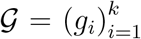 *be a sequence of gene trees such that for all* 1 *≤ i ≤ k, L*(*g*_*i*_) = *L, i*.*e*., *there is no missing data in any of the gene trees. If* |*L*| = *n, then there exists a pseudo quartet terrace of size at least* 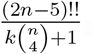.

*Proof*. Let *𝒯*_*L*_ be the set of all unrooted species trees on *L*. Note that we have |*𝒯*_*L*_| = (2*n −* 5)!!. There are a total of 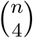 quartets in a tree with *n* taxa. Therefore, for a gene tree *g*_*i*_ *∈ 𝒢* and a species tree *T*, both on the same set of *n* taxa, *T* can satisfy at most 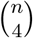 quartets (this is when *g*_*i*_ and *T* have an identical topology). Hence, for a set *𝒢* of *k* gene trees, the maximum number of consistent quartets with respect to a species tree *T* is 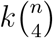. Therefore, we have 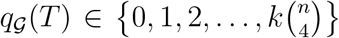. Now for 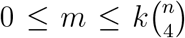, let *𝒯*_*m*_ = {*T ∈ 𝒯*_*L*_ : *q*_*𝒢*_ (*T*) = *m*}. Clearly, the sets *𝒯*_*m*_ form a partition of *𝒯*_*L*_ and by the pigeonhole principle, there exists *m* such that 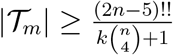. This set *𝒯*_*m*_ is our required pseudo quartet terrace.

Since the numerator in 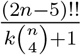 grows much faster with *n* than the denominator, the sizes of such pseudo terraces can grow very large very quickly even without missing data.

We also note that terraces – unlike pseudo terraces – are agnostic to gene tree topologies and depend solely on taxon coverage (i.e., the taxa sampled in gene trees). This is a consequence of Proposition 3.1 which says that if two trees display the same set of trees, then they have the same score for an additive scoring function. In other words, even if we replace each gene tree *g*_*i*_ with a different gene tree 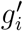, as long as the leaf sets remain the same, i.e. 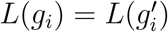, two trees *T*_1_ and *T*_2_ that were once in a terrace will remain in a terrace. So, in a very real sense, for species tree terraces, the topologies of the gene trees in the input sequence 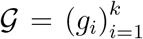 do not matter. The only thing that matters is the taxon coverage, i.e., the sequence of leaf sets 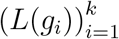. As long as this sequence of leaf sets in the gene trees is fixed, any two trees that are once in a species tree terrace with respect to *𝒢* will always remain in a species tree terrace for *𝒢*^*′*^ (albeit the optimality scores of the terraces could be different due to the topological differences of the gene trees in *𝒢* and *𝒢*^*′*^). Therefore, we have the following theorem.

### Theorem 3.2

*Let 𝒮 be a species tree terrace with respect to* 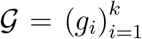 *of gene trees and an additive scoring function s* _*𝒢*_ (*·*). *Then 𝒮 is a species tree terrace with respect to every sequence* 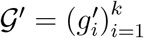 *where for* 1 *≤ i ≤ k*, 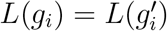.

Since the input gene tree topologies do not matter for determining whether a species tree belongs to a terrace, one set of species tree acts as a species tree terrace for an extremely large number of sequences of input gene trees as shown in the proposition below.

### Proposition 3.2

*Let* 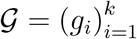 *be a sequence of gene trees and let 𝒮 be a species tree terrace with respect to 𝒢. Let ℳ*_*S*_ *be the set of all k gene tree sequences 𝒢*^*′*^ *for which 𝒮 is a species tree terrace. More formally*,

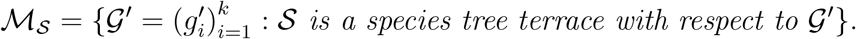

*Then* 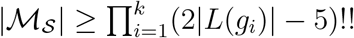.

*Proof*. Clearly, if 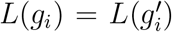 for all 1 *≤ i ≤ k*, then *𝒮* is a species tree terrace with respect to *𝒢*^*′*^. Since there are (2*n−*5)!! full binary trees on *n* leaves, the result follows.

The phylogenetic terrace, originally described by Sanderson *et al*. [21] in the context of estimating phylogenetic trees from sequence alignments, also depends solely on the taxon coverage. This is not a limiting factor for phylogenetic terraces as it is defined for a problem which takes sequence alignments as input, making the taxon coverage the only relevant information from the input data. Summary methods in the gene tree-species tree context, however, take a collection of gene trees – representing the taxon coverage of the input gene trees as well as gene tree topologies and their discordance. Taking the gene tree discordance into account is fundamental for estimating species trees from a collection of gene trees using a statistically consistent way. Therefore, it is desirable to have variants of species tree terraces that are sensitive to gene tree topologies in addition to the taxon coverage. In this regard, we introduce a special type of gene tree topology-aware terraces, which we call *peak terraces*.

## 4 Peak Terraces

Peak terraces are species tree terraces with one extra condition: for any tree *T* in a peak terrace, the sequence 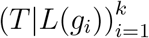 must equal *𝒢* (see Figure 4). More formally, we have the following definition.

**Figure 4:**
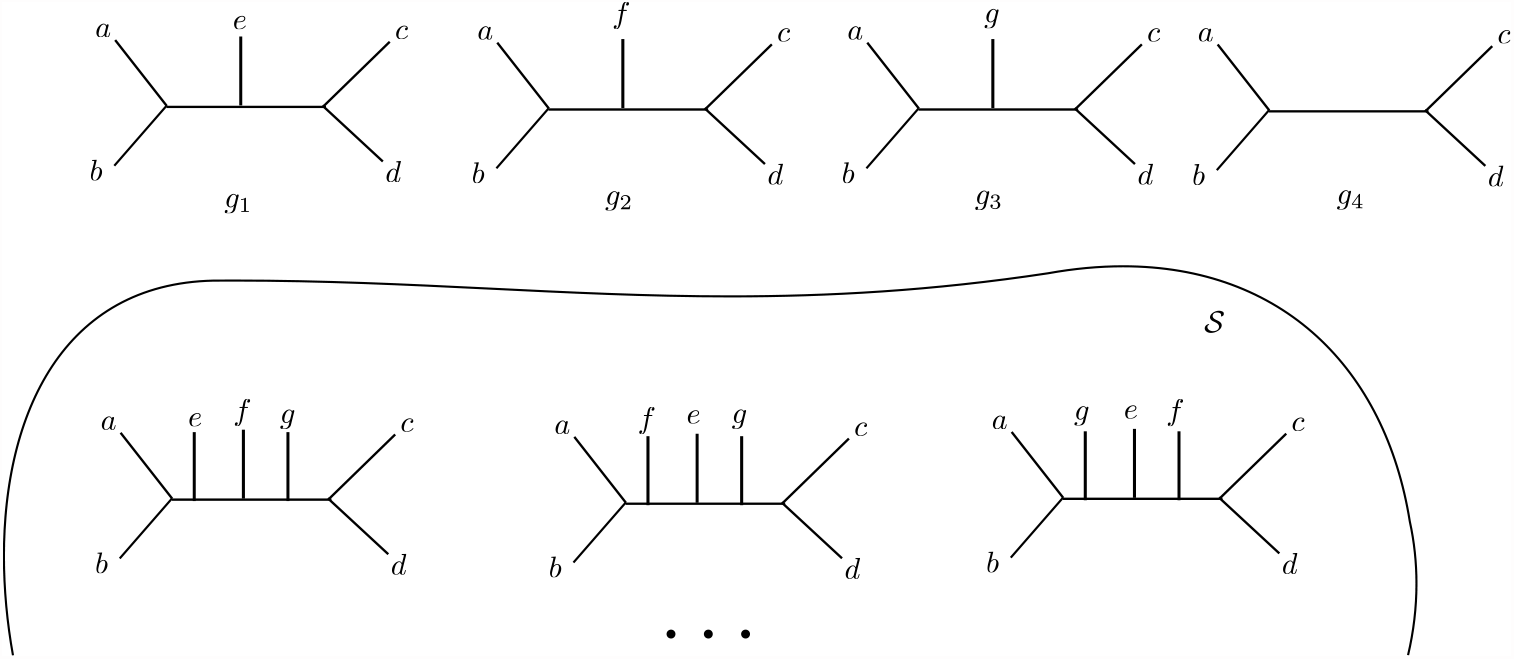
Portion of the peak terrace *𝒮* with respect to the sequence *𝒢* = (*g*_1_, *g*_2_, *g*_3_, *g*_4_) of gene trees. Every tree in *𝒮* displays each one of *g*_1_, *g*_2_, *g*_3_, and *g*_4_.

### Definition 4.1 (Peak Terrace)

*Let* 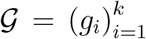 *be a sequence of gene trees. A set 𝒮 is called a peak terrace with respect to 𝒢, if for every T ∈ 𝒮*, 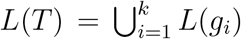 *and* 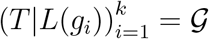.

Peak terraces are named as such due to the fact that any tree in a peak terrace achieves the optimal score with respect to the input sequence of gene trees. For maximization problems (e.g., maximizing quartet score), the trees in a peak terrace have the highest score, whereas for minimization criteria (e.g., minimizing deep coalescence, minimizing gene duplication and loss), the trees in a peak terrace achieve the lowest “cost” (i.e., the highest score, where score = *−*cost). We note that, in the landscape sense, the peak terraces for minimization problems are actually “basins” rather than “peaks”. The following proposition shows the optimality of the trees in a peak terrace in the context of quartet scores.

### Proposition 4.1

*Given a sequence* 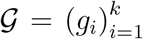 *of gene trees, any tree T in a quartet peak terrace with respect to 𝒢 must have the maximum possible quartet score with respect to 𝒢 i*.*e*., 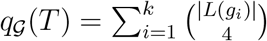.

*Proof*. Since *T* is in a quartet peak terrace, for every 1 *≤ i ≤ k, T* |*L*(*g*_*i*_) = *g*_*i*_. Therefore, *Q*(*T*) *⊇ Q*(*g*_*i*_), and so, 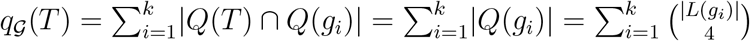. □

Note that Proposition 4.1 implies that for a fixed sequence *𝒢* of gene trees – unlike species tree terraces – there exists exactly one quartet peak terrace. However, we would like to point out that this is true for not only quartet peak terraces but also any *s*-terrace where *s* is an additive scoring function. More interestingly, for a particular sequence of gene trees, there is exactly one peak terrace for all possible additive scoring functions.

The concept of peak terraces has impacts on species tree inference from a collection of gene trees using summary methods. Summary methods attempt to find a tree with the optimal score (e.g., quartet score), meaning that when the problem is solved exactly, the output species tree will reside in a pseudo species terrace with the optimal score. The trees in such a terrace are all optimal in terms of an optimization criterion, but they are topologically different and hence have different topological accuracies – posing a challenge for the search algorithms to find comparatively reliable tree from a pool of equally optimal trees. This phenomenon was systematically analyzed and demonstrated by [26], where it was observed that Phylonet [34,35] (a method for estimating species trees by minimizing the number of extra lineages resulting from deep coalescence events) can produce trees with identical or competitive quartet scores as ASTRAL, but ASTRAL is typically substantially more accurate than Phylonet. Note that a pseudo species terrace with the optimal score is not necessarily a terrace or a peak terrace, but a peak terrace is a pseudo terrace with the optimal score, and both are sensitive to gene tree topologies. Moreover, as we will show in the following, characterizing peak terraces is sufficient for characterizing terraces. Understanding peak terraces may thus potentially help in the development of terrace-aware data structures and algorithms to circumvent the challenges and ambiguity posed by equally good trees, thereby improving tree search strategies for summary methods.

Another reason why peak terraces are important is the fact that every species tree terrace for some input sequence of gene trees is a peak terrace for possibly a different sequence of input gene trees. This different sequence is simply the sequence of gene trees displayed by the trees in the terrace when restricted to the leaves of the original sequence of input gene trees. In other words, we have the following fact.

### Fact 4.1

*Let* 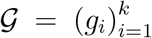 *be a sequence of gene trees. If 𝒮 is a non-empty species tree terrace with respect to 𝒢, then 𝒮 is also a peak terrace with respect to the sequence* 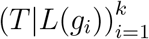 *where T ∈ 𝒮*.

Clearly, by definition, every peak terrace is a species tree terrace. Fact 4.1, however, tells us that every species tree terrace is also a peak terrace. It then follows that the set of all species tree terraces over all sequences of input gene trees is exactly the same as the set of all peak terraces over all sequences of input gene trees. So, if we want to understand the structural properties of terraces, it suffices to focus on the structural properties of peak terraces.

Although gene tree topologies do not matter for species tree terraces, they do matter for pseudo-terraces. Indeed, if one of the input gene trees is changed while keeping the leaf set unchanged, two trees that were once in a pseudo-terrace may cease to remain in it. The same is true for peak terraces too. Unlike species tree terraces which only depends on the leaf sets of the input gene trees, peak terraces are sensitive to the topologies of the gene trees. Moreover, it is not guaranteed that any species tree is a part of a peak terrace, meaning that the peak terrace can be necessarily empty for some set of inputs. The following fact highlights a simple sufficient condition for this to happen.

### Fact 4.2

*Let* 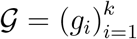 *be a sequence of gene trees such that there exists* 1 *≤ i < j ≤ k for which* 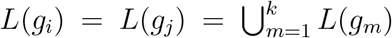 *but g*_*i*_ ≠ *g*_*j*_. *Then the peak terrace of 𝒢 is necessarily empty*.

Fact 4.2 tells us if there are two different gene trees that both contain all the taxa, then the peak terrace is necessarily empty. In other words, missing data is a necessity for peak terraces to be non-empty unless the input gene trees are all identical. However, that alone is not sufficient. The condition that all input gene trees be displayed is a rather strong one. A collection of trees is called *compatible* if there exists at least one tree that displays every tree in the collection. We have the following fact about peak terraces.

### Fact 4.3

*Let* 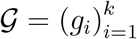 *be a sequence of gene trees. The peak terrace of 𝒢 is non-empty if and only if 𝒢 is compatible*.

Whether a collection of gene trees is compatible can be decided using the BUILD algorithm from [36].

### 4.1 Finding Patterns of Missing Data that Give Rise to Non-Trivial Peak Terraces

We now focus on the case where the input gene trees do not have any missing data. In other words, for each 1 *≤ i ≤ k*, 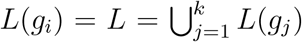. In this case, unless the input gene trees are not all identical, the peak terrace is necessarily empty. Therefore, we seek conditions on the patterns of missing data that cause the peak terrace to be non-empty. More formally, for 1 *≤ i ≤ k*, we seek *X*_*i*_ *⊆ L*(*g*_*i*_) with |*X*_*i*_| *≥* 4 such that there exists at least one tree *T* with leaf set 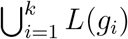 that displays each tree in the sequence 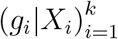. We call such a sequence 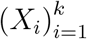 a *kernel*. Each kernel specifies a pattern of missing data for which a non-empty peak terrace exists.

We first show that if there are not too many gene trees in the input, then such a pattern of missing data can always be found.

#### Theorem 4.1

*Let* 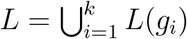 *be the set of all taxa present in the data. If the number of input gene trees, k, satisfies the inequality* 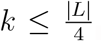, *then there exists a kernel i*.*e. a sequence* 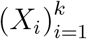 *with* |*X*_*i*_| *≥* 4 *such that there exists a T with L*(*T*) = *L that displays* 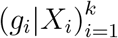.

*Proof*. Let *n* = |*L*| and let *L* = {*L*_1_, *L*_2_, …, *L*_*n*_}. For 1 *≤ i ≤ k*, we set *X*_*i*_ = {*L*_4*i−*3_, *L*_4*i−*2_, *L*_4*i−*1_, *L*_4*i*_} and let 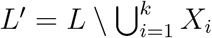. Note that for each 1 *≤ i ≤ k, g*_*i*_|*X*_*i*_ is a quartet. Finally, let *T* ^*′*^ be any full binary tree with leaf set *L*^*′*^. Now consider the tree *T* in Figure 5, which contains the quartets *g*_1_|*X*_1_, *g*_2_|*X*_2_, …, *g*_*k*_|*X*_*k*_, and the subtree *T* ^*′*^. Clearly, *T* displays 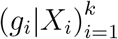.

**Figure 5:**
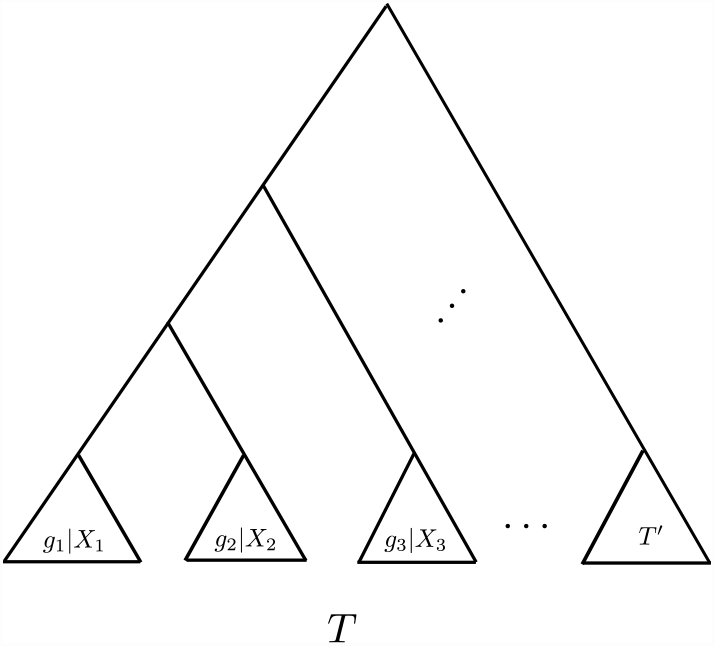
The tree *T* used in the proof of Proposition 4.1.

Theorem 4.1 gives a simple sufficient condition for the existence of a kernel based on the number of input gene trees. One might ask if there is an analogous necessary condition. It turns out there is not: no matter how many distinct gene trees are in the input sequence, there always exists a kernel provided that the gene trees have at least six leaves.

#### Theorem 4.2

*Let L be a set with* |*L*| *≥* 6 *and let* 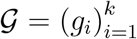 *be a sequence of gene trees such that for all* 1 *≤ i ≤ k, L*(*g*_*i*_) = *L. Then there exists a kernel i*.*e. a sequence* 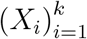 *with* |*X*_*i*_| *≥* 4 *such that there exists a tree T with L*(*T*) = *L that displays* 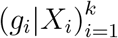.

To prove Theorem 4.2, we first need the following lemma.

#### Lemma 4.1

*Let L be a set such that* |*L*| = 6. *There exists a tree T with L*(*T*) = *L, such that for every tree T* ^*′*^ *with L*(*T* ^*′*^) = *L, Q*(*T*) *∩ Q*(*T* ^*′*^) ≠ *∅*.

*Proof*. Let *L* = {*a, b, c, d, e, f*}. We claim that the tree *T* in Figure 6 works.

**Figure 6:**
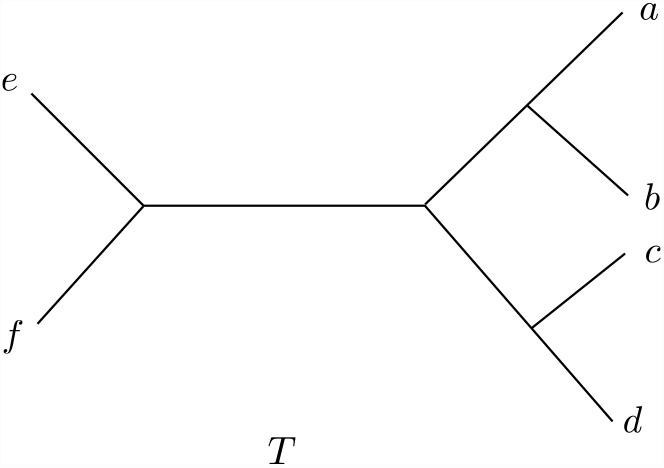
The tree *T* used in the proof of Lemma 4.1. For every tree *T* ^*′*^ with *L*(*T* ^*′*^) = *L*(*T*), *Q*(*T*) *∩ Q*(*T* ^*′*^) ≠ *∅*.

Assume, for the sake of contradiction, that there exists *T* ^*′*^ with *L*(*T* ^*′*^) = *L* such that *Q*(*T*) *∩ Q*(*T* ^*′*^) = *∅*. Then there exists a pair of leaves in *T* ^*′*^ that are siblings (i.e., the number of edges on the path between these two leaves is two). Without loss of generality, let *a* be one of these leaves and let *x* be its sibling. Note that *x* ≠ *b* since otherwise, every quartet of the form *ab* | *yz*, where *y, z ∈ {c, d, e, f*}, would be in *Q*(*T*) *∩ Q*(*T* ^*′*^). Let *x*^*′*^ be the sibling of *x* in *T*. Now choose leaves *y* and *z* from *L \ {a, x, b, x*^*′*^}. Clearly, since |*L*| = 6, this can always be done. Now the quartet *ax* | *yz ∈ Q*(*T*) *∩ Q*(*T* ^*′*^), a contradiction. □

Now we can proceed to prove Theorem 4.2.

*Proof of Theorem 4.2.* Let *L*^*′*^ *⊆ L* such that |*L*^*′*^| = 6. By Lemma 4.1, there exists a tree *T* ^*′*^ with *L*(*T* ^*′*^) = *L*^*′*^ such that *Q*(*L*^*′*^) *∩ Q*(*g*_*i*_|*L*^*′*^) ≠ *∅* for each 1 *≤ i ≤ k*. Now for each 1 *≤ i ≤ k*, choose *q*_*i*_ *∈ Q*(*L*^*′*^) *∩ Q*(*g*_*i*_|*L*^*′*^) and set *X*_*i*_ = *L*(*q*_*i*_). Clearly, *T* ^*′*^ displays 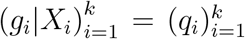, and so any tree *T* with *L*(*T*) = *L* that displays *T* ^*′*^ completes the proof. □

The bound in Theorem 4.2 is tight. If the gene trees all have fewer than six leaves, then for certain input sequences, it is possible for no kernel to exist. This is because the existence of a tree similar to the one in Lemma 4.1 is not guaranteed if |*L*| = 5.

#### Lemma 4.2

*Let L be a set such that* |*L*| = 5. *Then for every tree T with L*(*T*) = *L, there exists a tree T* ^*′*^ *with L*(*T* ^*′*^) = *L such that Q*(*T*) *∩ Q*(*T* ^*′*^) = *∅*.

*Proof*. Let *T* be any tree with *L*(*T*) = *L* and let *x ∈ L* have no sibling. Since |*L*| = 5, such an *x* can always be found. Let *q* = *T* |*L \ {x*} and *q*^*′*^ be a quartet on the same set of leaves as *q* (i.e., *L*(*q*^*′*^) = *L*(*q*) = *L \ {x*}) but is topologically different from *q*. Let *T* ^*′*^ be the tree obtained by attaching *x* to the internal branch of *q*^*′*^. It can be very easily seen that *Q*(*T*) *∩ Q*(*T* ^*′*^) = *∅*.□

Due to Lemma 4.2, if the input sequence 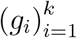 contains, say, all the gene trees on five leaves, then no kernel is possible. In other words, we have the following theorem.

#### Theorem 4.3

*Let X be a set with* |*X*| = 5. *If* 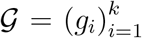 *is a sequence of gene trees containing every gene tree on X, then there does not exist a sequence* 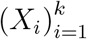 *with* |*X*_*i*_| *≥* 4 *such that there is a tree T*_1_ *that displays* 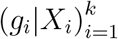.

*Proof*. For the sake of contradiction, assume there exists a sequence 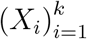 with |*X*_*i*_| *≥* 4 such that there is a tree *T*_1_ which displays 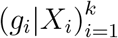. Let *T* = *T*_1_|*X*. Using Lemma 4.2, we can now obtain *T* ^*′*^ such that *L*(*T* ^*′*^) = *X* and *Q*(*T*) *∩ Q*(*T* ^*′*^) = *∅*. Since *G* contains every gene tree on *X*, it also contains *T* ^*′*^. And since *T* and *T* ^*′*^ have no quartets in common, neither do *T*_1_ and *T* ^*′*^. So, *T*_1_ can not display *T* ^*′*^, a contradiction. □

It is worth noting that even though Theorem 4.2 is stated in a way that disallows missing data, a similar result also holds when there is missing data as long as there is a set of at least six leaves common to every input gene tree.

#### Corollary 4.1

*Let* 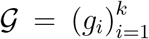 *be a sequence of (possibly incomplete) gene trees. If* 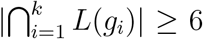, *then there exists* 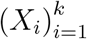 *with* |*X*_*i*_| *≥* 4 *such that there exists a T with* 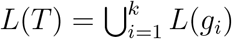 *that displays* 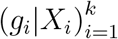.

*Proof*. We start with picking 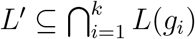 such that |*L*^*′*^| = 6. The rest of the proof is almost identical to that of Theorem 4.2. □

### 4.2 Extending to Other Scoring Functions

So far, we have only been focusing on quartet scores. For example, we wanted our *X*_*i*_’s to be of size at least four because according to our definition, the quartet score of a tree with fewer than four leaves is undefined. However, our results are general and can easily be extended to other additive scoring functions too. Consider, for example, the extra-lineage score for which we have the following analogous propositions. The proofs of these are left to the reader (a proof sketch for Theorem 4.4 is provided). Note that, unlike the quartet score, computing the extra-lineage score requires the gene trees and species trees to be rooted.

#### Proposition 4.2

*Given a sequence* 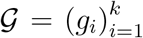 *of rooted gene trees, any tree T in an extra-lineage peak terrace with respect to 𝒢 must have an extra-lineage score of zero with respect to 𝒢*.

#### Theorem 4.4

*Let L be a set such that* |*L*| *≥* 6 *and let* 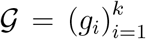 *be a sequence of rooted gene trees such that for all* 1 *≤ i ≤ k, L*(*g*_*i*_) = *L. Then there exists* 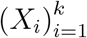 *with* |*X*_*i*_| *≥* 3 *for each i such that the extra-lineage peak terrace of* 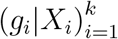 *is non-empty*.

*Proof sketch for Theorem 4*.*4*. The main idea is to find a *rooted* tree *T* such that for any rooted tree *T* ^*′*^ with *L*(*T*) = *L*(*T* ^*′*^), there exists at least one rooted triplet that is common to both *T* and *T* ^*′*^. It turns out that rooting the tree in Figure 6 on any branch results in a tree that works. Then we proceed similarly as the proof for Theorem 4.2.

## 5 Experimental results

We have designed the experimental study considering the following research questions (RQs).

- **RQ1:** Previous studies demonstrated the prevalence of pseudo terraces in phylogenomic analyses [37]. Species tree terraces in the context of estimating species trees from a collection of gene trees (as we have formalized in this study) have more stringent conditions for their existence as the trees in a terrace need to display the same locus-specific trees. As a result, how prevalent are species tree terraces under realistic model conditions, and what challenges do they present?
- **RQ2:** Can the accuracy of the estimated phylogenetic tree be enhanced through the utilization of species tree terraces?
- **RQ3:** Can we leverage trees within terraces to compute branch supports for the estimated species trees?
- **RQ4:** Do the true species trees reside in large terraces? Can we decrease such ambiguities by analyzing sufficiently large numbers of correct gene trees?

### Dataset description

We used the dataset generated and analyzed in [38], which presents a method ASTEROID for estimating species trees from gene trees in the presence of missing data. The data were generated using Simphy [39] by varying a wide range of parameters such as the level of missing data, the level of discordance due to ILS, the number of gene trees and the number of taxa. Gene trees were inferred using ParGenes [40] with one RAxML-NG maximum likelihood search from a single random starting tree per gene under the general time reversible model of nucleotide substitution with four discrete gamma rates (GTR+G4) [41,42]. Missing data was introduced by randomly sampling gene sequences, where each gene and each species has certain deletion probabilities (we refer to [38] for more details). The generated gene trees exhibit an average of around 60% missing taxa and 60% missing genes, making them suitable for investigating species tree terraces resulting from missing data. We also analyzed a biological dataset *Life92-single* with 92 species from the Eukaryote and Archaea domains [38, 43].

### Methods used

In order to examine the prevalance of species tree terraces considering various optimization criteria used for species tree construction (e.g., quartet score, RF score, etc.), we used a wide range of species tree estimation methods namely ASTRAL, wQFM, FastRFS, and ASTEROID.

### Measurements

To assess the quality of the estimated trees (on simulated datasets), we compared them with the model species tree using normalized Robinson-Foulds (RF) distance [44]. The RF distance between two trees is the sum of the bipartitions (splits) induced by one tree but not by the other, and vice versa. We also investigated the quartet scores (the number of quartets in the gene trees that agree with a species tree) of the trees estimated by different methods.

### 5.1 RQ1: Prevalence of species tree terraces

We first discover the terraces (if present) associated with the trees estimated by ASTRAL, which is arguably the leading coalescent-based method. Our experiments on 20 replicates of data containing 75 species and 1000 gene trees revealed that ASTRAL-estimated trees land on notably large terraces across all the replicates. For example, on a particular replicate (Rep-17) of this dataset, The tree estimated by ASTRAL-III [45] lands on a terrace of 14,175 trees. Since all of these 14,175 trees have the same quartet score, with respect to the gene trees under experimentation, any of these 14,175 trees could have been selected by ASTRAL. However, since these trees are topologically different, they have different RF scores with respect to the model species trees – raising an ambiguity for the tree search algorithm. To further investigate this, we calculated the RF scores of these 14,175 trees and plotted them against the corresponding quartet scores (Fig. 7a). Remarkably, despite having the same quartet score, the RF rates of these trees vary significantly, ranging from 0.236 - 0.347. Among the 14,175 trees within this particular terrace, we identified 707 trees that exhibited higher accuracy than the ASTRAL-estimated trees, 11,736 trees that were less accurate, and the remaining 1,732 trees displayed identical RF rates to the ASTRAL tree (in Fig. 7 and in subsequent discussion, these trees are denoted as “better”, “worse”, and “equal” trees respectively, each represented with a distinct colour).

**Figure 7:**
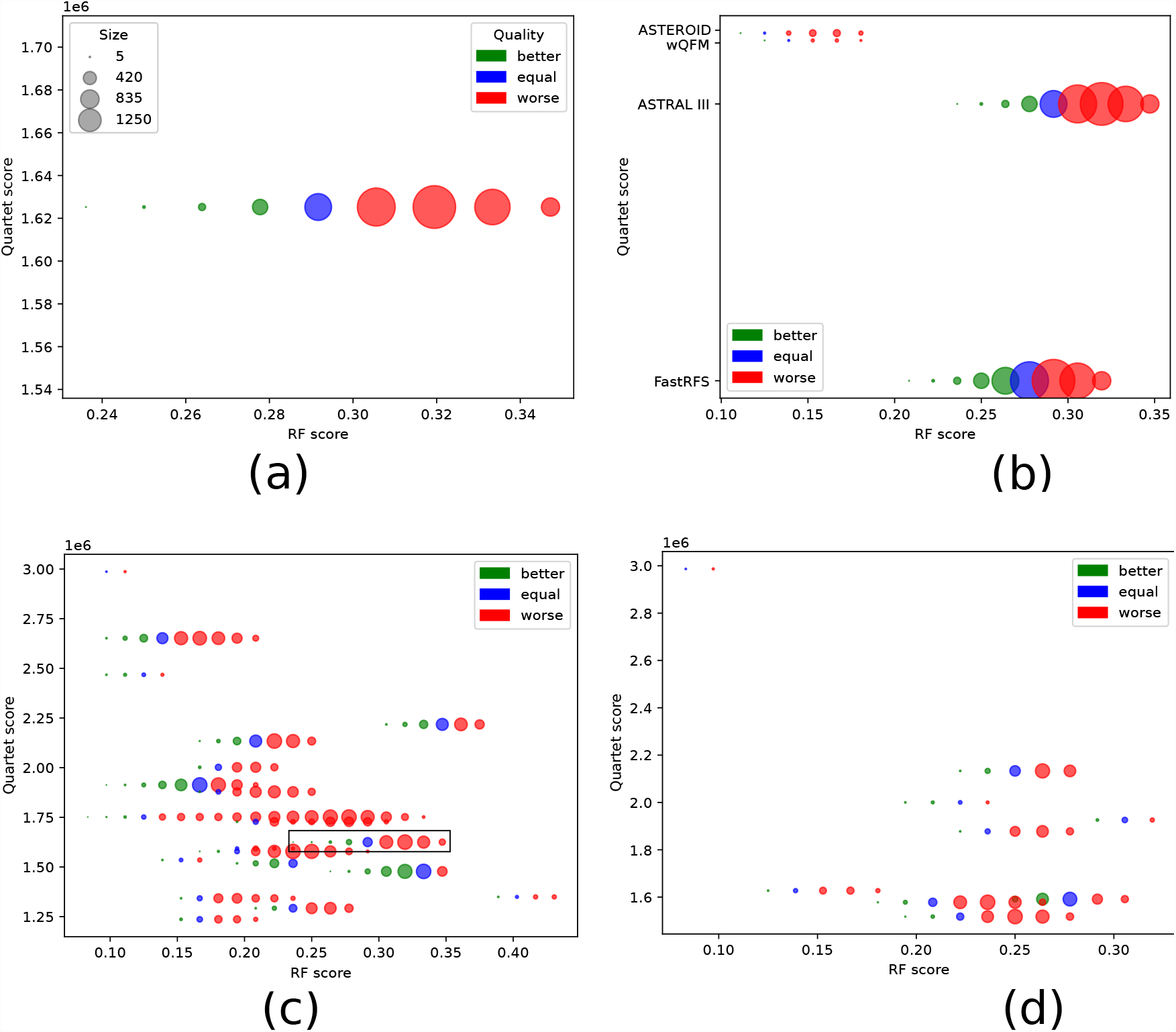
Prevalence of species tree terraces. (a) Quartet score vs. RF rate of the trees within the terrace, comprising 14,175 trees, which contains the tree estimated by ASTRAL on a particular replicate with 75 taxa. The better, worse, and equal trees (with respect to the ASTRAL-estimated tree) are shown in different colors. The size of the circles is proportional to the number of trees. (b) The species tree terraces corresponding to the trees estimated by ASTRAL, wQFM, ASTEROID, and FastRFS, and the variation in tree qualities of the trees within the terraces. (c) Presence of terraces across all replicates corresponding to the ASTRAL-estimated trees. Each horizontal line (represented by circles with different colors), positioned at different heights along the y-axis, represents a terrace corresponding to the trees estimated by ASTRAL on different replicates. The particular replicate that we show in Fig. 7a,b is marked by a rectangular box. (d) Presence of terraces across all replicates corresponding to the wQFM-estimated trees.

We then investigated if the prevalence of terraces and the variation of the tree qualities within the trees in the terraced landscape extend to other species tree estimation methods. Our investigations included the trees estimated by several popular methods, such as wQFM (which was shown to have superior accuracy compared to ASTRAL [18]), ASTEROID (specifically designed for handling missing data), and FastRFS. We found that the trees estimated by all these methods belong to terraces with different sizes and containing trees with diverse levels of tree accuracy and quartet scores (Fig. 7b). Each horizontal line Fig. 7b (represented by circles with different colors), positioned at different heights along the y-axis, represents a terrace corresponding to the trees estimated by different methods. Finally, we show in Fig. 7c that these observations hold across all the 20 replicates we examined for this dataset. Remarkably, better trees (than the tree estimated by ASTRAL) exist in the species tree terraces across all these 20 replicates, although their presence is limited in number for some of the replicates. In Fig. 7d, we demonstrate the same results as Fig. 7c but now we ran wQFM on the 20 replicates. Interestingly, we observed that the sizes of the terraces where the wQFM-estimated trees reside are substantially smaller than those of ASTRAL. Also, wQFM-estimated trees land on terraces with multiple species trees for 9 (out of 20) replicates, and no multiplicity of equally good trees was observed on the remaining 11 replicates.

Finally, we investigated the prevalence of terraces on other simulated datasets from [38] with varying numbers of taxa (25 - 125). The distributions of the trees inside the terraces across all these datasets are presented in Table 1. Similar trends, as observed on the 75-taxon dataset, were observed on other datasets. Note that the size of the terraces generally increases as we increase the number of taxa. This is expected because the number of candidate species trees grows exponentially with the number of taxa, making the terraces relatively large and thereby posing greater challenges to the tree search algorithms. wQFM-estimated trees generally belong to relatively small terraces while FastRFS-estimated trees land on larger terraces. As we mentioned earlier, this can be attributed to the relative accuracies of the trees estimated by different methods. Interestingly, we also observed that the number of better trees within the terrace is very small for wQFM compared to other methods, indicating that wQFM tends to select the tree with relatively high accuracy from within the corresponding terrace. In contrast, the terraces corresponding to other methods, such as FastRFS and ASTRAL, contain a substantial number of trees with better accuracy than the estimated trees. The underlying reasons for these different sizes of terraces for various methods remain unclear and warrant further investigation. It is possible that the diverse levels of terrace sizes are influenced by different algorithmic techniques and nuances employed to explore the search space during tree estimation.

**Table 1:** Average number of better, equal and worse trees in the terrace of the tree estimated by ASTRAL-III, ASTRAL-MP, FastRFS, wQFM, ASTEROID over 20 replicates of the 25-taxon to 125-taxon dataset.

| Number<br>of<br>Species | Method | Average<br>number of<br>better trees | Average<br>number of<br>equal trees | Average<br>number of<br>worse trees | Average<br>Total Number<br>of trees |
| --- | --- | --- | --- | --- | --- |
| 25 | ASTRAL III | 89.8 | 110.6 | 1,384.9 | 1,585.3 |
|  | FastRFS | 12.2 | 30.2 | 1,028.4 | 1,070.8 |
|  | wQFM | 0.5 | 3.7 | 25.0 | 29.2 |
|  | Asteroid | 12.0 | 25.6 | 1,364.1 | 1,401.6 |
| 50 | ASTRAL III | 397.9 | 413.2 | 1,245.7 | 2,056.8 |
|  | FastRFS | 550.2 | 410.9 | 1,306.3 | 2,267.3 |
|  | wQFM | 647.4 | 638.8 | 494.6 | 1,780.8 |
|  | Asteroid | 74.7 | 198.7 | 1,130.1 | 1,403.4 |
| 75 | ASTRAL III | 191.0 | 244.4 | 2,146.8 | 2,582.2 |
|  | FastRFS | 195.7 | 305.6 | 1,777.6 | 2,278.8 |
|  | wQFM | 14.2 | 31.7 | 155.6 | 201.4 |
|  | Asteroid | 122.0 | 154.9 | 1,379.8 | 1,656.7 |
| 100 | ASTRAL III | 780.1 | 823.6 | 5,664.8 | 7,268.4 |
|  | FastRFS | 978.9 | 915.6 | 5,565.7 | 7,460.1 |
|  | wQFM | 30.0 | 69.4 | 443.6 | 543.0 |
|  | Asteroid | 1,687.2 | 1,182.1 | 3,894.8 | 6,764.1 |
| 125 | ASTRAL III | 778.3 | 952.3 | 7,035.8 | 8,766.3 |
|  | FastRFS | 1,395.6 | 1,176.3 | 7,120.0 | 9,691.8 |
|  | wQFM | 4.8 | 21.8 | 2,659.6 | 2,686.2 |
|  | Asteroid | 351.0 | 536.0 | 5,907.8 | 6,794.7 |

#### Prevalence of terraces in biological dataset

We analyzed the *Life92-single* dataset with 3199 single-copy gene trees covering 92 species from the Eukaryote and Archaea domains [38, 43]. Morel *et al*. [38] compiled these single-copy gene trees from 41,222 multicopy gene trees originally inferred by Williams *et al*. [43] by applying DISCO and filtering out resulting single-copy gene trees with less than four species.

We found that the species tree estimated by ASTRAL from these gene trees belongs to a very vast terrace with an astonishing number of 9,38,81,025 (around 94 million) trees, and all with an identical quartet score of 4,04,36,626. Interestingly, the ASTEROID-estimated tree lands on a separate terrace but has the same number of trees (9,38,81,025), yielding a quartet score of 4,03,23,847.

These experiments demonstrate that species tree terraces, despite having a stringent condition of displaying identical locus-specific trees, may frequently occur in phylogenomic studies. More importantly, while the presence of terraces introduces ambiguity and challenges for the tree search algorithms, it presents opportunities for finding more accurate trees. Our findings demonstrate that, in almost all cases, some trees within these terraces are better than the trees estimated by existing methods. For example, the average RF rate of ASTRAL-estimated trees over 20 replicates of the 25-taxon dataset is 0.285 while the average RF rate of the best trees within the terraces containing ASTRAL-estiamted trees is 0.189, showing that there exist substantially better trees than ASTRAL despite having identical quartet scores. Therefore, in the next section, we investigate the potential for leveraging the trees in a species tree terrace for finding relatively accurate species trees.

### 5.2 RQ2: Leveraging terraces for improved phylogenomic analysis

In Sec 5.1, we observed that within a terrace containing a tree estimated by a specific method (e.g., ASTRAL, wQFM, etc.), there are more accurate trees than the estimated ones. Hence, it becomes crucial to identify these relatively accurate trees from within the terrace. However, as discussed earlier in Proposition 3.1 and Corollary 3.1, all trees in a terrace display identical locus-specific trees and thus share the same scores based on any additive optimization criteria (e.g., quartet score, triplet score, extra lineage score, duplication/loss score, etc.). Consequently, distinguishing relatively accurate trees based on optimization criteria alone is not feasible.

One approach could involve computing consensus trees of the trees within a terrace. However, considering that there are both better and less accurate trees (than the estimated ones) in a terrace, computing consensus trees of all the trees may not yield improved accuracy. In contrast, our experiments demonstrate that computing the consensus of only the better trees significantly enhances results compared to computing the consensus of all trees within the terrace.

In Fig. 8 a, b, for each method (e.g., ASTRAL, wQFM, etc.), we show the average RF rates of the following four types of trees: i) trees estimated by a particular method, ii) consensus trees of the trees within the terrace that contains the trees estimated by that particular method, iii) consensus trees of only the better trees from within the terrace, and iv) the best tree, in terms of the tree accuracy, within the terrace containing the estimated tree. The results demonstrate that computing the consensus of all trees within the terrace is not beneficial, as it combines the less accurate trees (which often outnumber the relatively good trees), leading to a decline in overall performance. On the other hand, computing the consensus of only the better trees yields significant improvements for all methods across all datasets. Notably, as shown in Fig. 8, the accuracies of these consensus trees are comparable to the best trees in the terraces. This highlights the potential for enhancing species tree accuracy by identifying a subset of the terrace that contains more trees with better accuracy than worse trees. However, effectively distinguishing such trees remains a challenge that requires further investigation in future studies. By addressing this challenge, we can potentially improve species tree estimation in the presence of terraced landscapes.

**Figure 8:**
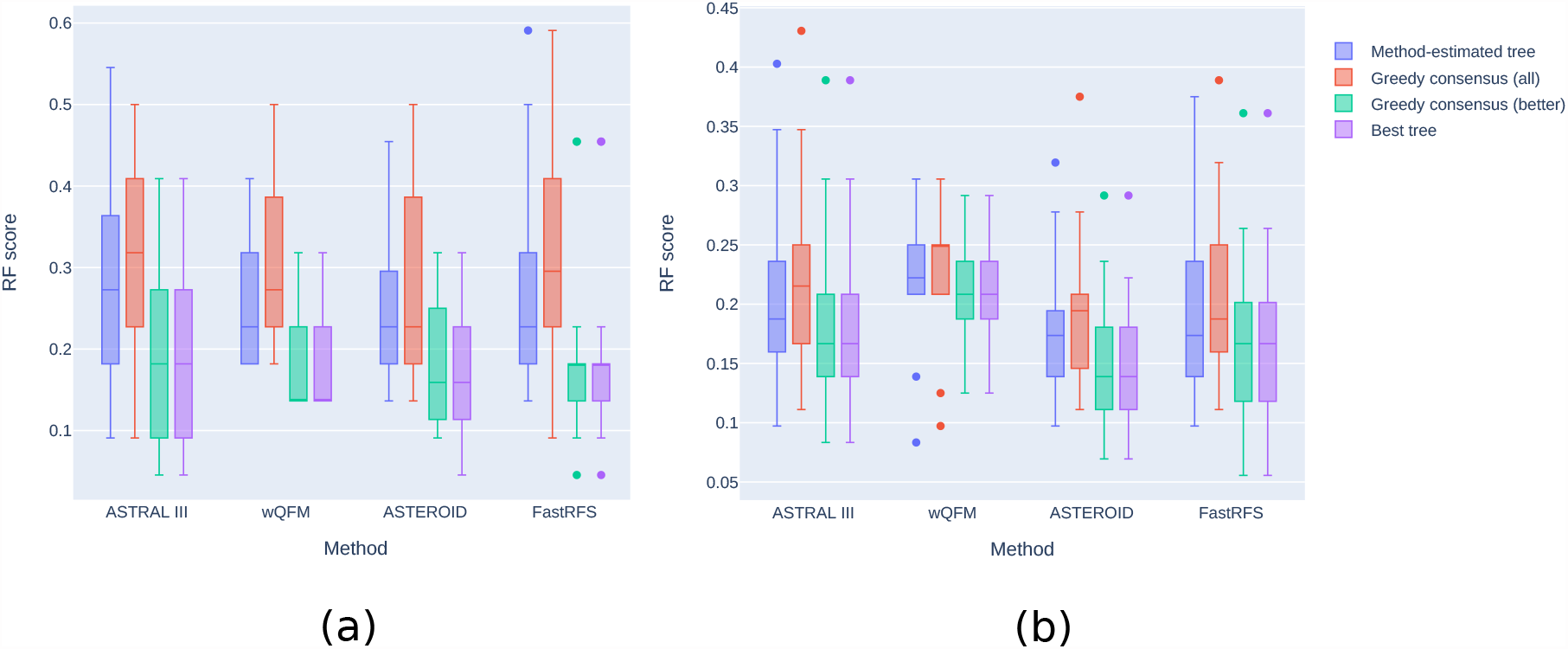
Leveraging terraces for improved phylogenomic analysis. We show the average RF rates (over 20 replicates) of different methods and compare them to the greedy consensus trees computed from the trees within their respective terraces. Consensus trees were estimated from all the trees as well as from only the better trees within each terrace. Additionally, the average RF rate of the best tree within a terrace is presented. Results are shown for both the 25-taxon dataset (a) and the 75-taxon dataset (b).

### 5.3 RQ3: Leveraging terraces for inferring branch supports

We infer branch supports on an estimated tree based on the trees within the terrace that contains the estimated tree. Support on a specific branch was computed based on the fraction of trees (out of trees within the terrace) that contain that specific branch (i.e., bipartition). We compared the branch support estimated using this way to the supports computed using local posterior probabilities [46] computed by ASTRAL. We assess the quality of the branch supports using the following two metrics.

#### Calibration

We first bin branches by their support into several groups and quantify the relationship between bins of branch support and the percentage of correctly placed queries in each bin. For example, for branches in the 50-60% support bin, we say the results are calibrated if roughly 55% of these branches are correct.

#### Empirical Cumulative Distribution Function (ECDF)

Support values can be analysed by studying their ECDF, which involves separating the accurate and inaccurate branches. Ideally, incorrect branches have low support (uniformly distributed) and correct branches have high support (depending on the signal, and hence, the power). Generally, a wider difference between the distribution of correct and incorrect branches is desired.

The support values obtained by both terrace-based estimation and ASTRAL are well calibrated with the accuracy (i.e., support values are closer to the expected values) as shown in Fig. 9 a, b. The difference between ASTRAL- and terrace-based methods are more pronounced at lower support levels, where the terrace-based method tends to underestimate support values. For branches with relatively high accuracy, the terrace-based method tends to assign higher supports than ASTRAL, as evidenced by the larger dot denoting *∼*100% support compared to the support inferred by ASTRAL.

**Figure 9:**
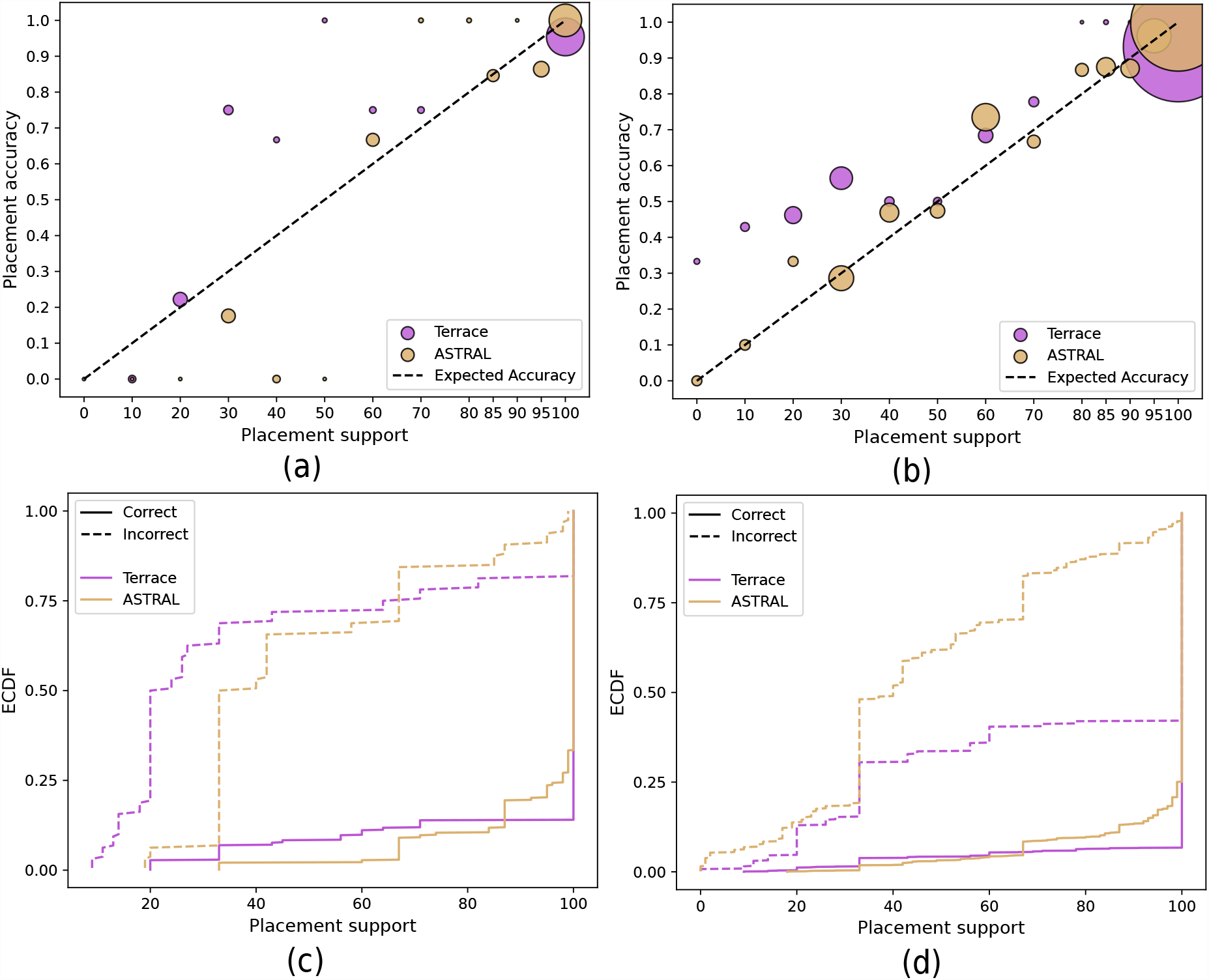
Comparison of terrace-based supports to the supports computed using local posterior probabilities inferred by ASTRAL. (a)-(b) Support versus the percentage of correctly placed queries over twenty replicates of data on 25-taxon and 75-taxon datasets, respectively. Support values are binned at 0%, 10% … 80%, 85%, 90%, 95%, and 100% left inclusive (e.g., [0,10)); the last bin only includes 100%. Unity line (*y* = *x*): fully-calibrated support. Dot sizes are proportional to the number of branches within each bin. (c)-(d) Empirical cumulative distribution function (ECDF) of the support for correct and incorrect placements for 25- and 75-taxon datasets, respectively.

The results can be better understood by examining the Mean Squared Errors (MSEs) of accuracy (computed with respect to the unity line) as presented in Table 2. For branches with higher support values (*≥* 80%), terrace-based supports have comparable MSEs to those of ASTRAL, with a difference that is almost negligible. However, terrace-based support exhibits limited effectiveness in low support ranges (*<* 80%) with notably higher MSEs compared to ASTRAL-based support values.

**Table 2:** The mean squared error (MSE) of the points computed with respect to the unity line and shown separately for low (*<* 80%) and high (*≥*80%) support values. We compare terrace-based supports and ASTRAL-based supports.

| Number of Species | Low Support |  | High Support |  |
| --- | --- | --- | --- | --- |
|  | Terrace-based | ASTRAL-based | Terrace-based | ASTRAL-based |
| 25 | 1.712 | 1.204 | 0.007 | 0.002 |
| 50 | 6.141 | 3.526 | 0.003 | 0.001 |
| 75 | 8.265 | 2.103 | 0.005 | 0 |
| 100 | 22.675 | 4.221 | 0.006 | 0.001 |
| 125 | 18.372 | 1.883 | 0.003 | 0 |

Investigating the distribution of the support values, there is a large gap between the support distribution of correct and incorrect placements. For 25-taxon dataset, the gap is more prominent for terrace-based supports than the ASTRAL-based supports. With terrace-based supports, 50% of the incorrect branches have less than 20% support whereas 50% of the incorrect branches have less than 35% support with ASTRAL-based method, showing the superiority of terrace-based approach. However, as we increase the number of taxa, the support values for incorrect branches estimated by ASTRAL becomes more meaningful than terrace-based methods as it assigns higher proportion of incorrect branches with relatively low support values. For correct branches, both terrace- and ASTRAL-based approaches perform well, with the majority of correct branches receiving 100% support. Notably, the terrace-based approach demonstrates a slight advantage over ASTRAL on the correct branches, as evidenced by the corresponding ECDF curve dipping below the one corresponding to ASTRAL-based support. This indicates that the terrace-based approach assigns more accurate branches with 100% support.

Overall, despite the existence of substantial numbers of less accurate trees in terraces, terrace-based approach was able to infer well-calibrated and meaningful support values, showcasing the promise of using the trees within a terrace for inferring support values.

### 5.4 RQ4: Investigating the terraces containing true species trees and the impact of increasing numbers of correct input gene trees on terraced landscape

We have already demonstrated the presence of terraces and the associated ambiguity for species trees estimated by different summary methods. We now investigate if the true species trees also belong to large terraces. Indeed, as we show in Fig. 10, true species trees are also contained within substantially large species tree terraces. This further emphasizes the uncertainty arising from terraces, as it becomes impossible to distinguish the trees within a terrace based on any additive optimization criteria. Consequently, finding the true tree using summary methods under practical model conditions, even when they are statistically consistent, remains uncertain. This could be attributed to the fact that the number of input gene trees is limited and there is estimation error in them. Given a sufficiently large number of correct gene trees (i.e., no estimation error), summary methods that optimize statistically consistent measures such as quartet and triplet scores will converge in probability to the true species tree. That means, if we increase the number true gene trees, the size of the terraces containing the true species tree should gradually decrease to one, allowing for the unique identification of the true species tree using statistically consistent methods.

**Figure 10:**
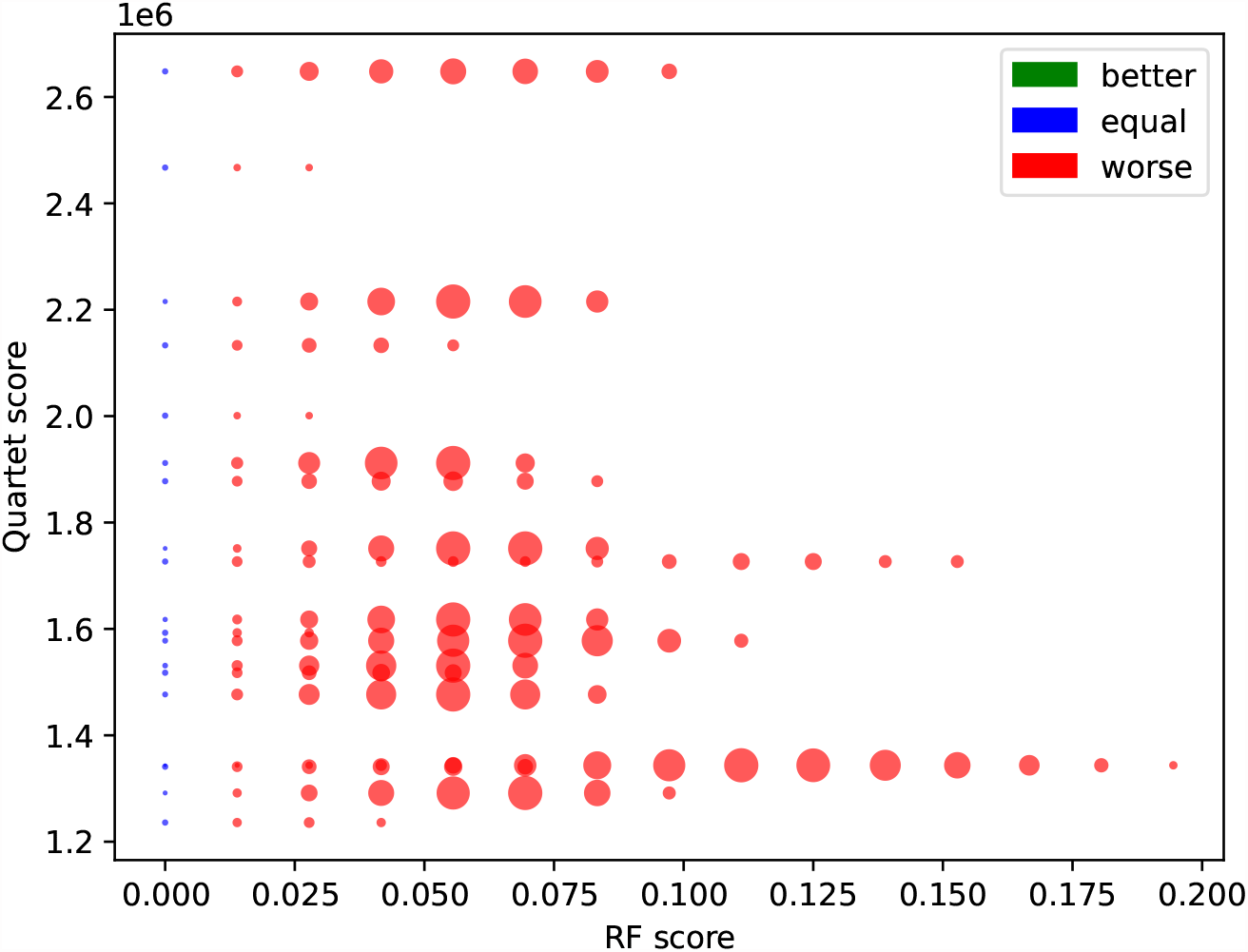
Distribution of species tree terraces that contain the true trees. We show quartet score versus RF rate of the trees within the species tree terrace, which contain the tree true species tree across all the replicates of the 75-taxon dataset. 19 out of 20 replicates demonstrate the presence of species tree terraces containing the true species tree. The true species tree always has an RF rate of 0, resulting in the blue line along the y-axis at an RF score of 0. As there exists no better tree than the true species tree, green dots are absent here. The sizes of the terraces across different replicates vary substantially and thus we normalize the size of the terraces for better visualization. As a result, the relative size of species tree terraces is not conserved.

To further investigate this, we explore the size of terraces containing the true species tree in relation to the number of true gene trees. Fig. 11 presents the results on two representative replicates from the 25-taxon dataset. As expected, the size of the species tree terraces corresponding to the true tree asymptotically decreases with increasing numbers of true gene trees. Eventually, the size of the terraces should gradually decrease to one – containing only the true species tree. Simultaneously, the quartet score increases with an increasing number of true gene trees and decreasing sizes of the terraces. These asymptotic trends suggest that, with sufficiently large number of true gene trees, the quartet score is likely to reach its maximum when a terrace contains only one tree, which is the true species tree. This indicates that maximizing statistically consistent measures like the quartet score enables the unique identification of the true species tree with a high probability when a sufficiently large number of true gene trees is available.

**Figure 11:**
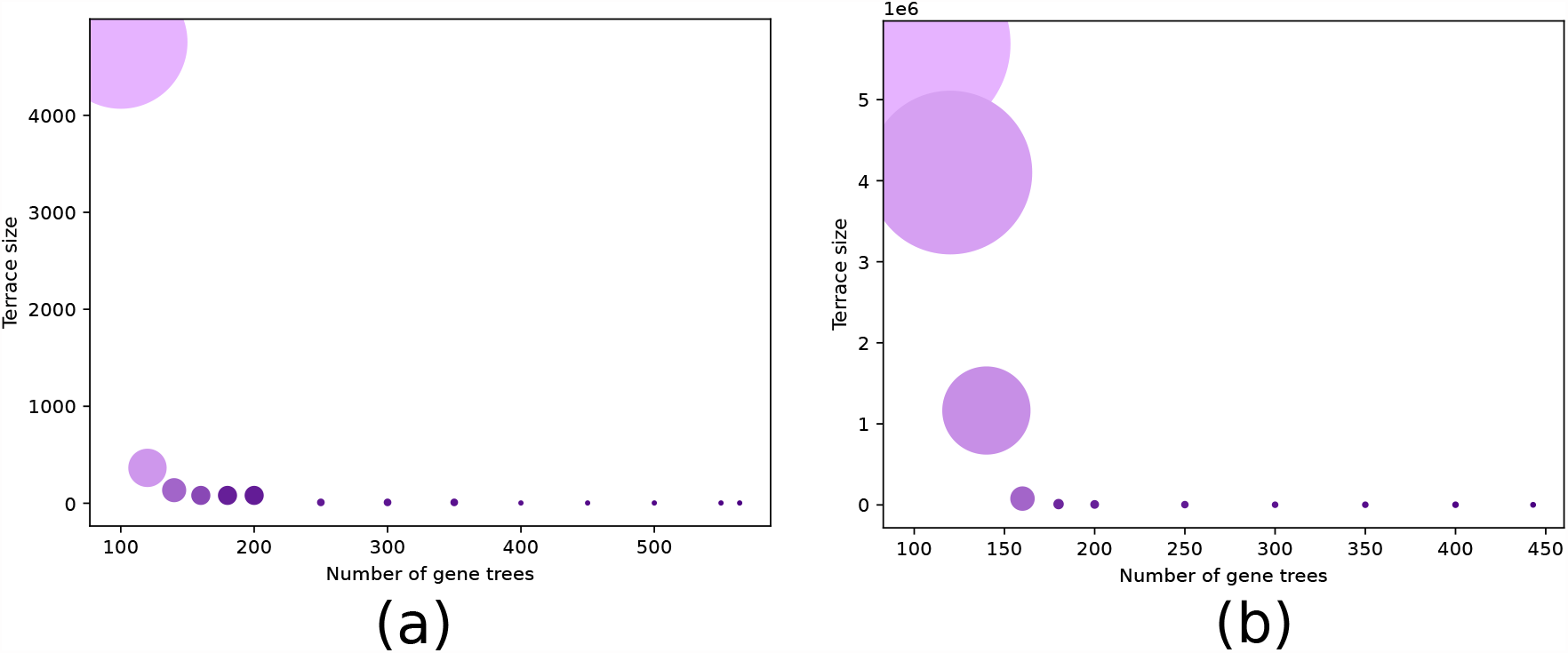
Size of terraces in relation to the number of true gene trees. We vary the number of true gene trees from 100 to 564 and, for each case, show the size of the terraces. We present the results on two replicates of data, shown separately in (a) and (b), from the 25-taxon dataset. The size of a circle is proportional to the number of trees within a terrace. We color the circles with a color gradient which varies continuously from light purple to dark purple with increasing quartet scores.

## 6 Conclusions

The mathematical characterizations of species tree terraces and the new results on terraces in the gene tree-species tree context presented here are timely and important as large genome-scale phylogenomic studies with substantial amounts of missing data are becoming increasingly common [47, 48]. The multiplicity of equally good species trees in terraced landscapes, stemming from certain patterns of missing data, poses various challenges as well as opens up several important research avenues. The ability to detect if a tree resides on a terrace, computing the size of the terrace, and enumerating the trees in a terrace can potentially improve the scalability and accuracy of summary methods. However, detecting and exploring species tree terraces, and leveraging that to improve the tree search strategies are challenging. Discovering various combinatorial properties of terraces and conditions for the presence of multiple equally good trees can contribute towards developing terrace-aware data structures and tree search algorithms.

We formally characterized species tree terraces and contrasted them with pseudo species tree terraces. We showed that, unlike pseudo terraces, species tree terraces depend only on the taxon coverage and are agnostic about the gene tree topology. However, considering different genes having different topologies is central to developing statistically consistent species tree estimation methods. In this study, we introduced a new type of terrace called peak terrace which requires one additional condition than terraces, making it sensitive to the distribution of gene tree topologies. Moreover, despite the differences in their mathematical definitions and conditions for existence, we showed that the set of species terraces is identical to the set of peak terraces. Therefore, we argued that understanding the structural properties of peak terraces suffices to understand species tree terraces in general. We proved various combinatorial properties of peak terraces and investigated patterns of missing data that lead to the existence of peak terraces. Although we explicitly considered the quartet score for investigating some of the properties of terraces, our results are general and apply to other additive scoring functions. We systematically performed a set of experiments investigating the presence of species tree terraces and the associated challenges and opportunities. We found that substantially more accurate species trees compared to the estimated ones can be found from within terraces if we are able to distinguish them using appropriate optimization criteria. Thus, we believe that this study will prompt more analytical and experimental studies to better understand the terraced landscapes of species trees and pioneer new terrace-aware methods, data structures, and optimization criteria for computing species trees from gene trees despite missing data and gene tree heterogeneity.

This study can be extended in several directions. Efficient terrace-aware data structures and algorithms for systematically navigating trees both inside a species tree terrace and its neighborhood would contribute to the improvement of the summary methods both in terms of accuracy and scalability. Therefore, investigating how to adapt the summary methods and algorithms to the existence of terraces is one of the most interesting research avenues. Developing efficient tools to identify species tree terraces and enumerating trees in them (similar to the existing tools [49] for counting trees in phylogenetic terraces) for different optimality criteria (e.g., quartet score, extra lineage score, etc.) is another important research direction. There are many questions of theoretical interest as well. For example, Section 4.1 considers the problem of finding the existence of a kernel given a gene tree sequence. A natural extension to this would be to find a kernel of the maximum size. Such a kernel would maximize the sum of the number of leaves remaining in the gene trees. One might also try to generalize Lemma 4.1, which asserts the existence of a binary tree *T* on six leaves whose quartets intersect every other binary tree on the same leaf set. One way to generalize this would be to consider trees larger than quartets. In other words, given *k*, let *n*(*k*) be the smallest number for which there exists a binary tree *T* on *n*(*k*) leaves such that for every binary tree *T* ^*′*^ with *L*(*T* ^*′*^) = *L*(*T*), there is at least one *k*-leaf binary tree that is displayed by both *T* and *T* ^*′*^. By Lemma 4.1, we have *n*(4) = 6. One might be interested in *n*(*k*) for *k >* 4 and ask how it grows with *k*. Overall, future studies need to investigate further combinatorial properties of species tree terraces, the challenges they pose, and solutions to these problems.

## Data Availability

The datasets analyzed in this study are available at https://cme.h-its.org/exelixis/material/asteroid_data.tar.gz.

## References

[1] W. P. Maddison. Gene trees in species trees. Systematic Biology, 46:523–536, 1997.

[2] Sebastien Roch and Mike Steel. Likelihood-based tree reconstruction on a concatenation of aligned sequence data sets can be statistically inconsistent. Theoretical Population Biology, 100:56–62, 2015.

[3] L S Kubatko and J H Degnan. Inconsistency of phylogenetic estimates from concatenated data under coalescence. Systematic Biology, 56:17, 2007.

[4] Scott V Edwards, Liang Liu, and Dennis K Pearl. High-resolution species trees without concatenation. Proceedings of the National Academy of Sciences, 104(14):5936–5941, 2007.

[5] A D Leaché and B Rannala. The accuracy of species tree estimation under simulation: a comparison of methods. Systematic Biology, 60(2):126–137, 2011.

[6] Michael DeGiorgio and James H Degnan. Fast and consistent estimation of species trees using supermatrix rooted triples. Molecular Biology and Evolution, 27(3):552–569, 2009.

[7] Md Shamsuzzoha Bayzid and Tandy Warnow. Naive binning improves phylogenomic analyses. Bioinformatics, 29(18):2277–2284, 2013.

[8] J Heled and A J Drummond. Bayesian inference of species trees from multilocus data. Molecular Biology and Evolution, 27:570–580, 2010.

[9] E Mossel and S Roch. Incomplete lineage sorting: consistent phylogeny estimation from multiple loci. IEEE/ACM Transactions on Computational Biology and Bioinformatics, 7(1):166–171, 2011.

[10] L S Kubatko, B C Carstens, and L L Knowles. Stem: Species tree estimation using maximum likelihood for gene trees under coalescence. Bioinformatics, 25:971–973, 2009.

[11] Siavash Mirarab, Rezwana Reaz, Md S Bayzid Théo Zimmermann, M Shel Swenson, and Tandy Warnow. ASTRAL: genome-scale coalescent-based species tree estimation. Bioinformatics, 30(17):i541–i548, 2014.

[12] Liang Liu, Lili Yu, and Scott V Edwards. A maximum pseudo-likelihood approach for estimating species trees under the coalescent model. BMC Evolutionary Biology, 10:302, 2010.

[13] Liang Liu and Lili Yu. Estimating species trees from unrooted gene trees. Systematic Biology, 60(5):661–667, 2011.

[14] B Larget, S K Kotha, C N Dewey, and C Ané. BUCKy: Gene tree/species tree reconciliation with the Bayesian concordance analysis. Bioinformatics, 26(22):2910–2911, 2010.

[15] David Bryant, Remco Bouckaert, Joseph Felsenstein, Noah A Rosenberg, and Arindam RoyChoudhury. Inferring species trees directly from biallelic genetic markers: bypassing gene trees in a full coalescent analysis. Molecular Biology and Evolution, 29(8):1917–1932, 2012.

[16] Julia Chifman and Laura Kubatko. Quartet from SNP data under the coalescent model. Bioinformatics, 30(23):3317–3324, 2014.

[17] Mazharul Islam, Kowshika Sarker, Trisha Das, Rezwana Reaz, and Md Shamsuzzoha Bayzid. Stelar: A statistically consistent coalescent-based species tree estimation method by maximizing triplet consistency. BMC Genomics, 21(1):1–13, 2020.

[18] Mahim Mahbub, Zahin Wahab, Rezwana Reaz, M Saifur Rahman, and Md Shamsuzzoha Bayzid. wQFM: highly accurate genome-scale species tree estimation from weighted quartets. Bioinformatics, 37(21):3734–3743, 2021.

[19] Rezwana Reaz, Md Shamsuzzoha Bayzid, and M Sohel Rahman. Accurate phylogenetic tree reconstruction from quartets: A heuristic approach. PLoS ONE, 9(8):e104008, 2014.

[20] Y. Yu, T. Warnow, and L. Nakhleh. Algorithms for MDC-based multi-locus phylogeny inference: Beyond rooted binary gene trees on single alleles. Journal of Computational Biology, 18(11):1543–1559, 2011.

[21] Michael J Sanderson, Michelle M McMahon, and Mike Steel. Terraces in phylogenetic tree space. Science, 333(6041):448–450, 2011.

[22] Michael J Sanderson, Michelle M McMahon, Alexandros Stamatakis, Derrick J Zwickl, and Mike Steel. Impacts of terraces on phylogenetic inference. Systematic biology, 64(5):709–726, 2015.

[23] Olga Chernomor, Arndt Von Haeseler, and Bui Quang Minh. Terrace aware data structure for phylogenomic inference from supermatrices. Systematic Biology, 65(6):997–1008, 2016.

[24] Katherine St. John. The shape of phylogenetic treespace. Systematic Biology, 66(1):e83–e94, 2017.

[25] Barbara H Dobrin, Derrick J Zwickl, and Michael J Sanderson. The prevalence of terraced treescapes in analyses of phylogenetic data sets. BMC Evolutionary Biology, 18(1):46, 2018.

[26] Ishrat Tanzila Farah, Muktadirul Islam, Kazi Tasnim Zinat, Atif Hasan Rahman, and Shamsuzzoha Bayzid. Species tree estimation from gene trees by minimizing deep coalescence and maximizing quartet consistency: a comparative study and the presence of pseudo species tree terraces. Systematic Biology, 70(6):1213–1231, 2021.

[27] Alexandros Stamatakis and Michael Ott. Efficient computation of the phylogenetic likelihood function on multi-gene alignments and multi-core architectures. Philosophical Transactions of the Royal Society B: Biological Sciences, 363(1512):3977–3984, 2008.

[28] Alexandros Stamatakis and Nikolaos Alachiotis. Time and memory efficient likelihood-based tree searches on phylogenomic alignments with missing data. Bioinformatics, 26(12):i132–i139, 2010.

[29] Lam-Tung Nguyen, Heiko A Schmidt, Arndt Von Haeseler, and Bui Quang Minh. Iq-tree: a fast and effective stochastic algorithm for estimating maximum-likelihood phylogenies. Molecular Biology and Evolution, 32(1):268–274, 2015.

[30] Michael J Sanderson, Michelle M McMahon, and Mike Steel. Terraces in gene tree reconciliation-based species tree inference. bioRxiv, 2020.

[31] L. Zhang. From gene trees to species trees II: Species tree inference by minimizing deep coalescence events. IEEE/ACM Transactions on Computational Biology and Bioinformatics, 8(9):1685–1691, 2011.

[32] M. S. Bayzid, S. Mirarab, and T. Warnow. Inferring optimal species trees under gene duplication and loss. In Proc. of Pacific Symposium on Biocomputing (PSB), volume 18, pages 250–261, 2013.

[33] M. S. Bayzid and T. Warnow. Gene tree parsimony for incomplete gene trees: addressing true biological loss. Algorithms for Molecular Biology, 13:1, 2018.

[34] C. V. Than, D. Ruths, and L. Nakhleh. PhyloNet: A software package for analyzing and reconstructing reticulate evolutionary relationships. BMC Bioinformatics, 9:322, 2008.

[35] Yun Yu, Tandy Warnow, and Luay Nakhleh. Algorithms for mdc-based multi-locus phylogeny inference: beyond rooted binary gene trees on single alleles. Journal of Computational Biology, 18(11):1543–1559, 2011.

[36] Alfred V. Aho, Yehoshua Sagiv, Thomas G. Szymanski, and Jeffrey D. Ullman. Inferring a tree from lowest common ancestors with an application to the optimization of relational expressions. SIAM Journal on Computing, 10(3):405–421, 1981.

[37] Ishrat Tanzila Farah, Md Muktadirul Islam, Kazi Tasnim Zinat, Atif Hasan Rahman, and Md Shamsuzzoha Bayzid. Phylogenomic terraces: presence and implication in species tree estimation from gene trees. bioRxiv, 2020.

[38] Benoit Morel, Tom A Williams, and Alexandros Stamatakis. Asteroid: a new algorithm to infer species trees from gene trees under high proportions of missing data. Bioinformatics, 39(1):btac832, 2023.

[39] Diego Mallo, Leonardo de Oliveira Martins, and David Posada. Simphy: phylogenomic simulation of gene, locus, and species trees. Systematic biology, 65(2):334–344, 2016.

[40] Benoit Morel, Alexey M Kozlov, and Alexandros Stamatakis. Pargenes: a tool for massively parallel model selection and phylogenetic tree inference on thousands of genes. Bioinformatics, 35(10):1771–1773, 2019.

[41] Simon Tavaré. Some probabilistic and statistical problems on the analysis of dna sequence. Lecture of Mathematics for Life Science, 17:57, 1986.

[42] Ziheng Yang. Maximum-likelihood estimation of phylogeny from dna sequences when substitution rates differ over sites. Molecular biology and evolution, 10(6):1396–1401, 1993.

[43] Tom A Williams, Cymon J Cox, Peter G Foster Gergely J Szöllősi, and T Martin Embley. Phylogenomics provides robust support for a two-domains tree of life. Nature ecology & evolution, 4(1):138–147, 2020.

[44] D.F. Robinson and L.R. Foulds. Comparison of phylogenetic trees. Mathematical Biosciences, 53:131–147, 1981.

[45] Chao Zhang, Maryam Rabiee, Erfan Sayyari, and Siavash Mirarab. Astral-iii: polynomial time species tree reconstruction from partially resolved gene trees. BMC Bioinformatics, 19(6):153, 2018.

[46] Erfan Sayyari and Siavash Mirarab. Fast coalescent-based computation of local branch support from quartet frequencies. Molecular Biology and Evolution, 33(7):1654–1668, 2016.

[47] Erich D Jarvis, Siavash Mirarab, Andre J Aberer, Bo Li, Peter Houde, Cai Li, Simon YW Ho, Brant C Faircloth, Benoit Nabholz, Jason T Howard, et al. Wholegenome analyses resolve early branches in the tree of life of modern birds. Science, 346(6215):1320–1331, 2014.

[48] Norman J Wickett, Siavash Mirarab, Nam Nguyen, Tandy Warnow, Eric Carpenter, Naim Matasci, Saravanaraj Ayyampalayam, Michael S Barker, J Gordon Burleigh, Matthew A Gitzendanner, et al. Phylotranscriptomic analysis of the origin and early diversification of land plants. Proceedings of the National Academy of Sciences, 111(45):E4859–E4868, 2014.

[49] Rudolf Biczok, Peter Bozsoky, Peter Eisenmann, Johannes Ernst, Tobias Ribizel, Fedor Scholz, Axel Trefzer, Florian Weber, Michael Hamann, and Alexandros Stamatakis. Two c++ libraries for counting trees on a phylogenetic terrace. Bioinformatics, 34(19):3399–3401, 2018.

